# RpiBeh: a multi-purpose open-source solution for real-time tracking and behavior-driven closed-loop interventions in rodent neuroethology

**DOI:** 10.1101/2024.12.02.626497

**Authors:** Yiqi Sun, Jie Zhang, Qianyun Wang, Jianguang Ni

## Abstract

High-precision behavior tracking and closed-loop intervention are essential for studying the neural basis of cognition and behavior. Existing commercial systems are costly and inflexible for customization, while current open-source tools are often lack of real-time functionality and suffer from steep learning curve. To address these issues, we developed RpiBeh, an open-source, cost-effective, and versatile software tailored for rodent neuroethological research. The software features an intuitive interface with extensive customization options. RpiBeh leverages a Raspberry Pi and camera for video streaming, enabling behavior-driven closed-loop control. Additionally, it provides frame-by-frame video timestamp output for precise synchronization with external devices. For real-time tracking and locomotion pattern analysis, RpiBeh utilizes several novel algorithms and integrated newly developed deep-learning method. Specifically, we introduced two algorithms: a Background Subtraction Method (BSM) for real-time position tracking and a Frame Difference (FD) algorithm for freezing behavior detection. RpiBeh was validated in single animal real-time tracking and locomotion pattern detection, demonstrating flexibility and effectiveness in configurating behavior-triggered closed-loop reinforcement experiments including passive place avoidance task and social fear conditioning tasks. It achieved the same level of performance in tracking and locomotion pattern detection comparing to benchmark software including ANY-maze and DeepLabCut, with superior customization and expandability. Consequently, RpiBeh offers an efficient, affordable, and open-source solution for video tracking and behavior-driven closed-loop experiments.

## 1. Introduction

In modern neuroscience research, high-precision behavior tracking and intervention is critical for causally interrogating the neural underpinnings of cognition and behavior. Such systems enable real-time readout of posture and locomotion information, providing the basis for temporally precise regulation and reinforcement of complicated behavior. Depending on how intervention is triggered, these systems can be categorized as closed-loop or open-loop systems. In recent years, behavior-driven (e.g., spatial region-of-interesting, posture, social interaction) closed-loop intervention has demonstrated substantial advantages in studying rodent navigation, decision-making, and social cognition ^1–3^, which often outperformed traditional open-loop experiments in terms of efficiency, specificity, and personalization ^1,4^.

Despite its promising potentials in neuroethology research, closed-loop experiments demand real-time processing of data streaming, posing challenges for its effective and user-friendly implementation. Although some existing commercial systems have offered variated solutions, these platforms often come with high costs and limited expandability. In recent years, open-source platforms have emerged as popular alternatives in behavioral monitoring ^5,6^, pose estimation ^7–15^ and data streaming control ^16,17^. Yet, many of these solutions only have limited real-time applicability, lacking of precise synchronization between behavior and external devices, and may present steep learning curve.

To address these gaps, we developed RpiBeh, a python based, affordable multi-purpose platform for rodent neuroethological research. It supports multiple common behavioral paradigms (e.g., social behavior, learning and memory task, open-field test, etc.), offers a user-friendly interface, and retains rich customization capabilities. The system uses a Raspberry Pi & camera for video streaming to support behavior-dependent closed-loop control. Moreover, it supports high-level frame-by-frame output of video timestamps, enabling precise synchronization between video and other external device (e.g., Open Ephys ^18^). For real-time tracking and locomotion analysis, the software utilizes several novel algorithms and integrated newly developed deep-learning methods (e.g., DLC-Live ^19^) to secure robust performance comparable to popular commercial software and mainstream open-source solutions.

We demonstrated the functionality and performance of RpiBeh in serval mice learning and social tasks. In a passive place avoidance task (PPA) ^20^, mice were reinforced to avoid a punishment zone by 2D spatial region of interesting (spatial-ROI) triggered mild electric foot shocks. In a social fear conditioning task (SFC) ^3^, a classic model for studying social anxiety disorders, the testing mouse received electric foot shocks when approaching a conspecific social mate, causing social avoidance behavior. The sound performance of RpiBeh together with its cost-effective and good expandability underscores its potential wide application in rodent neuroethology.

## 2. Materials and methods

### 2.1 System Design

We present an open-source software platform designed for real-time analysis and closed-loop intervention of mice behavior, which integrates with a Raspberry Pi (Raspberry Pi Ltd, UK) to facilitate video recording and supports a variety of behavioral experiments at affordable cost.

The hardware setup for video recording is depicted in Fig.1A. The system consists of a control host responsible for software control and analytical processing, and a Raspberry Pi for video capture and GPIO signal output. Communication between the control host and the Raspberry Pi is established either through Ethernet or wireless network. The high-level video control policy in the Raspberry Pi has been adapted from ^5^, enabling it to receive commands from the host via the ZeroMQ (ZMQ) protocol. During video capture, each frame is transmitted to the control host in real-time. Additionally, the system is capable of generating a brief 1-millisecond 3.3-volt GPIO signal at each frame capture, as shown in Fig.1B, thereby enabling precise synchronization with other external devices.

**Fig. 1.**
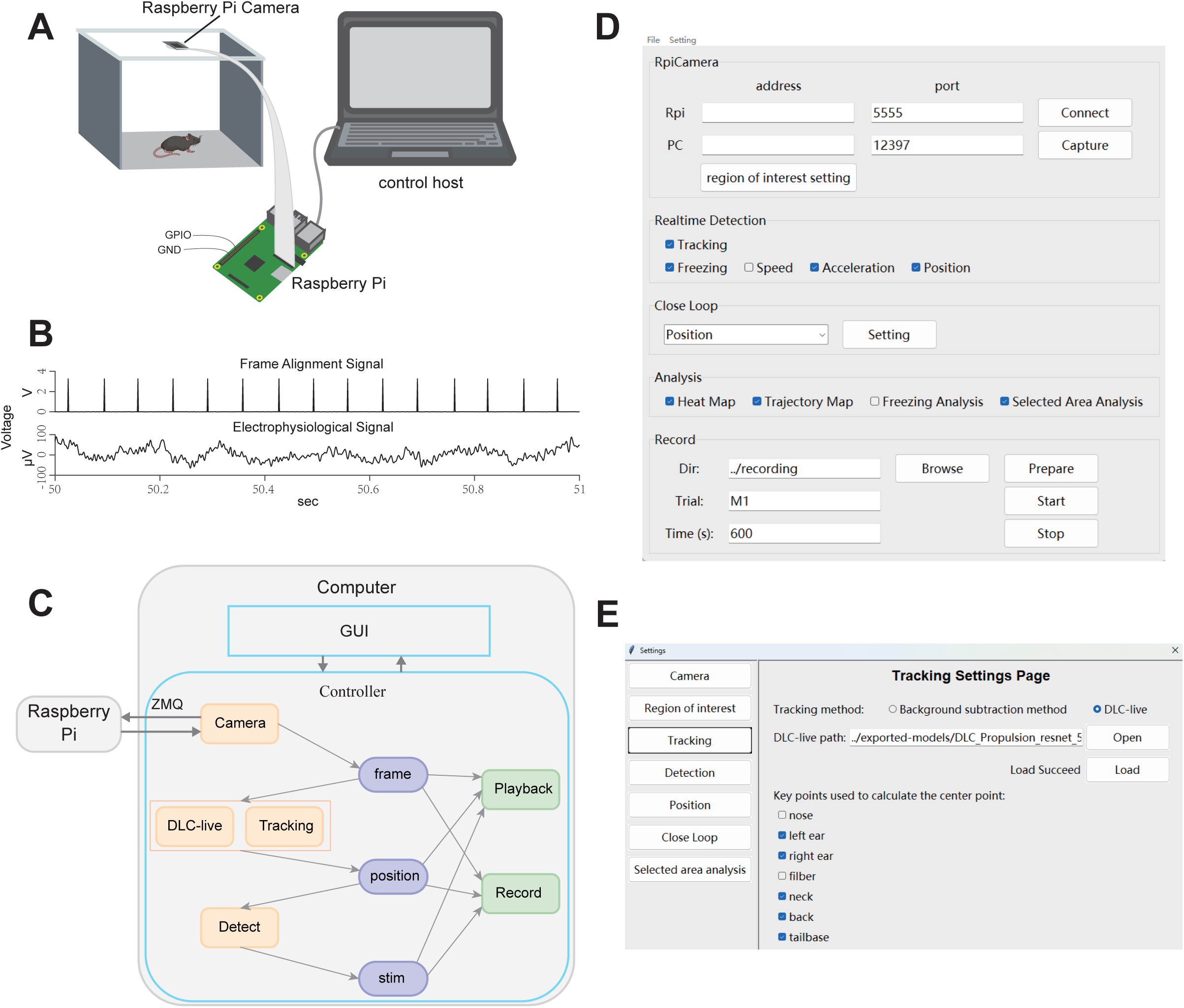
System design of the RpiBeh platform. (A). Schematic diagram of hardware including the Raspberry Pi, Raspberry Pi Camera, and the control host. Closed-loop control signals is transmitted through GPIO wiring to connected external devices. (B). Example of synchronizing video frame timestamps with electrophysiological recordings. (C). Schematic diagram the software architecture of the RpiBeh. (D). Example of the RpiBeh GUI. (E). The RpiBeh parameter settings page in the GUI.

The software’s design architecture is shown in Fig.1C. A graphical user interface (GUI) facilitates user interaction and control of the experiment, while internally the system is composed of multiple modules, including a camera module, a tracking module, various real-time detection modules, playback, and recording modules. These modules operate independently using a multithreaded architecture, with buffered data transmission between them to ensure that delays in any one module do not affect the overall performance.

The overall closed-loop control workflow is as follows: the Raspberry Pi camera captures video data stream and transmits it in frames to the control host in real-time. The control host detects the mouse’s position in each frame and performs state assessments (e.g., freezing or entry into specified ROI) based on locomotion data. The system then generates a stimulation command, which is send out through the Raspberry Pi’s GPIO interface to achieve targeted control.

The GUI for RpiBeh is illustrated in Fig.1D. Users input the IP addresses and ports for both the Raspberry Pi and the control host, then initiate connection via the “Connect” button, and capture a background image with the “Capture” button. The software enables users to specify real-time detection metrics (e.g., mouse position, freezing state, speed, acceleration, or entry into manually specified 2D spatial ROI) and customize the selected closed-loop control options. Recordings can be initiated for a specific duration and can be manually terminated at any time using the “Stop” button.

As shown in Fig.1E, the software supports a variety of configuration settings. Users can adjust video recording parameters such as frame rate and resolution, define the spatial ROI, set scale factors, and select tracking methods (currently supporting both the DLC-Live method and the Background Subtraction Method). Additionally, users can configure detection parameters such as threshold and duration, designate detection zones for ROI-based entry tracking, and define behavior-driven closed-loop control parameters (e.g., stimulus duration and frequency). The system also allows for post hoc analysis, such as calculating and summarizing locomotion distance, duration of stay, and the number of entries within defined ROI.

### 2.2 Tracking Methods

The software provides two methods for mouse position tracking: DLC-Live ^19^ and Background Subtraction Method (BSM).

#### DLC-Live Method

When using the DLC-Live method, a customized DeepLabCut ^9^ network must first be trained and exported. The network model is then imported through the software’s settings page by specifying its path. After successful import, users can select the desired key points from a provided list to compute the mouse’s center position. The center position for each frame is calculated by averaging the coordinates of the selected key points.

Due to the complexity of network training and the dependence of deep learning computations on hardware performance, the real-time efficiency of the DLC-Live method can be limited, especially on systems without a GPU. Therefore, the software also offers the Background Subtraction Method as an alternative for scenarios with constrained computing resources.

#### Background Subtraction Method

The workflow of the Background Subtraction Method (BSM) is illustrated in Fig.2. Before video recording begins, users are prompted to capture a background image without the presence of the mouse. During recording, each frame undergoes the following steps to determine the mouse’s center of mass (CoM):

1. Background Subtraction: Subtract the background image from the current frame to remove static elements from the scene;
2. Thresholding: Apply thresholding to the difference image to convert it into a binary format, isolating the mouse as the foreground;
3. Morphological Operations: Use morphological opening to remove noise from the foreground, followed by morphological closing to fill small holes, ensuring the continuity of the mouse’s region;
4. Connected Component Analysis: Identify and extract the largest connected component (the mouse), and calculate its centroid to represent the mouse’s position.

**Fig. 2.**
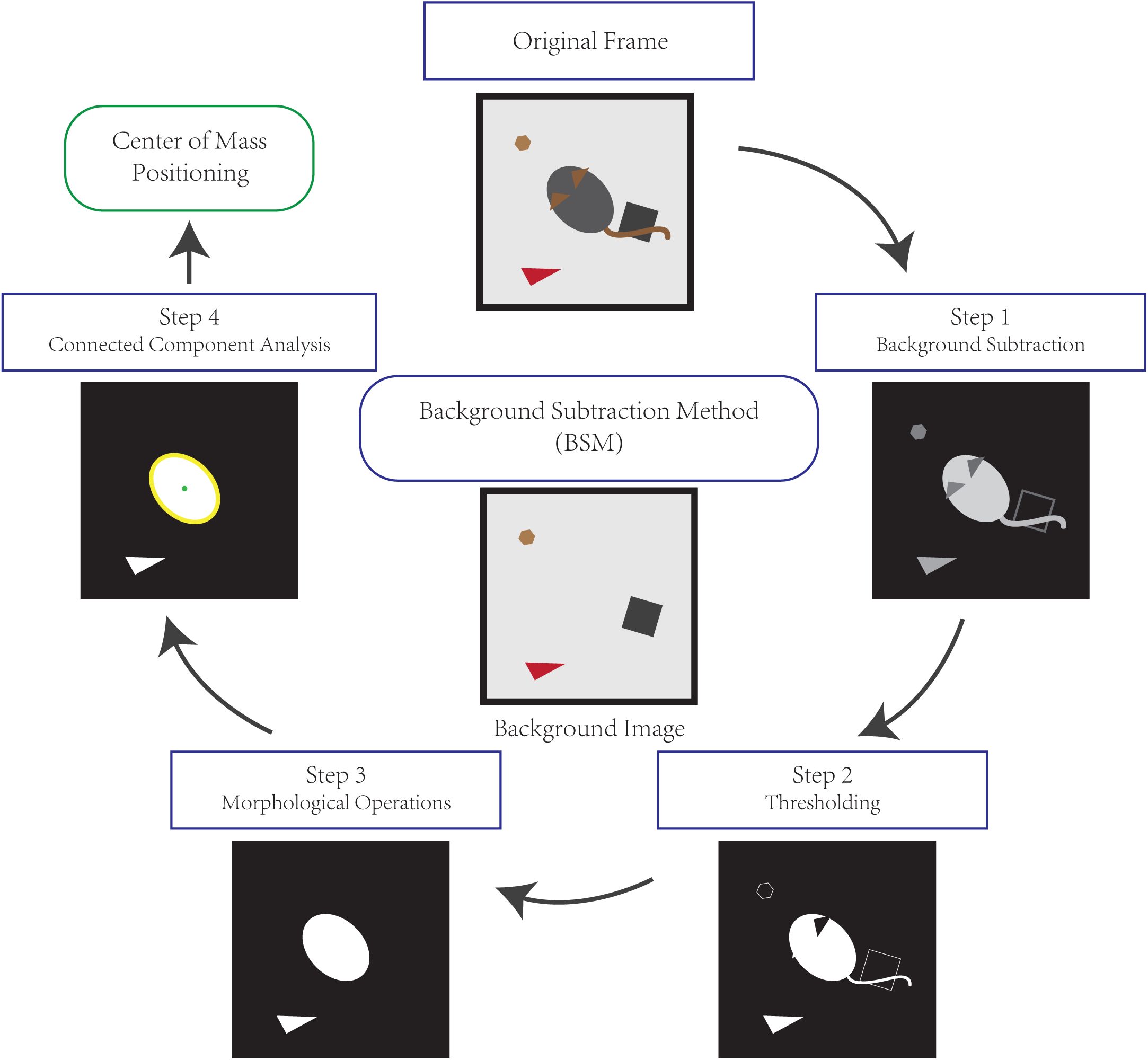
General procedure of the BSM for mouse position tracking. Positioning of the center of mass is determined by four steps: background subtraction, thresholding, morphological operations and connected component analysis.

This design balances flexibility and computational efficiency, providing a reliable solution for mouse tracking across various experimental conditions.

### 2.3 Online locomotion pattern identification

#### Online Freezing Detection

This experiment employs the Frame Difference (FD) algorithm to detect freezing state of mice in Pavlovian contextual fear conditioning task. Before recording, users need to configure a freezing threshold and a duration threshold in the software’s settings. The detection of the freezing state is determined by whether the frame nonmatching ratio between two consecutive frames falls below the threshold for a specified duration. The detailed steps of the algorithm are as follows:

1. Frame Difference: Frame difference calculation involves subtracting the pixel values of the preceding frame from current frame. To normalize the difference and avoid division by zero, each pixel value of the difference is divided by the corresponding pixel value in the preceding frame with a small constant (i.e., 5) was added. The result is then scaled to a range of 0–255. This ensures numerical stability while preserving the relative differences between frames.
2. Binarization: Apply binarization to the difference image to distinguish between pixels with and without changes. First, the difference image is separated into three channels of primary colors (Red, Green, and Blue). Each channel has a pixel intensity range of 0 to 255. A threshold value of 120 is selected for binarization, meaning that pixels with intensity values greater than or equal to 120 are set to 1 (or white), while those below 120 are set to 0 (or black). This binarization process is applied independently to each of the three channels. Afterward, a logical AND operation is performed across the binarized results of the three channels to generate a final binarized grayscale difference image.
3. Frame Nonmatching Ratio: Compute the percentage of all pixels that has value of one from the binarized grayscale difference image.
4. Freezing State Detection: Compare the frame nonmatching ratio with the predefined threshold. If the difference rate remains below the threshold for the specified duration, the mouse is identified as freezing.

During real-time detection, certain frames may satisfy the threshold criteria but fall short of the required duration. As a result, these frames cannot be identified as freezing states in real time, potentially causing misclassification. To address this limitation, an offline FD algorithm is employed to refine the detection. After the recording is completed, the offline algorithm re-analyzes all frames with pixel difference rates below the threshold and marks them as fear states if they maintain the condition for the specified duration, ensuring more accurate detection.

After the experiment, the RpiBeh provides a frame nonmatching ratio curve to illustrate the results of the freezing state analysis, offering a visual representation of the mouse’s behavioral states. Before conducting the formal experiment, users can perform a short trial recording to adjust parameters and select appropriate threshold values based on these visual outputs.

Finally, manually annotated freezing states served as the ground truth for evaluating the accuracy of automated freezing detection methods. The freezing states were labeled by human observers and subsequently reviewed by an independent researcher to ensure reliability.

#### Online Locomotion Speed Detection

The locomotion speed is calculated using the CoM obtained in real-time from the tracking process. For each pair of consecutive frames, the distance moved by the CoM is divided by the time interval between the two frames to compute the instantaneous speed. To reduce noise and enhance the robustness of the speed measurement, a moving average of 0.5 seconds is applied to the calculated speeds. Based on the speed threshold and duration requirements predefined by the experimenter, real-time speed detection results are provided.

### 2.4 Benchmarking algorithms for tracking and freezing detection

To evaluate the performance of RpiBeh, we compared its online tracking and freezing detection capabilities with several widely used methods, including ANY-maze, DeepLabCut (DLC), and an accelerometry-based approach. Each method was tested under controlled conditions using pre-recorded videos that mimicked real-time operations, ensuring that the performance of the algorithms was consistent with how they would operate during live video analysis.

#### RpiBeh Online Detection

For the analysis, pre-recorded videos were processed in a way that simulated real-time video streaming, leveraging code that mimicked the Raspberry Pi’s video capture and streaming process using ZeroMQ (ZMQ). This setup ensured that the RpiBeh algorithms—specifically for mouse tracking and behavioral detection—operated under the same conditions as in live experiments. The video frames were streamed in real-time from locally stored files, maintaining consistency between offline analysis and real-time performance.

#### ANY-maze Online Detection

ANY-maze was employed to track, record, and analyze the location and fear behavior of the mice. This commercially available video tracking software is widely used in research, known for its recognized accuracy and reliability, making it a standard benchmark for behavioral analysis.

#### DeepLabCut Offline Detection and DLC-Live Online Tracking

We employed the DeepLabCut (DLC) (8) system for mouse tracking and freezing detection. Seven key points (nose, left ear, right ear, fiber, neck, back, and tail base) were annotated, and the ResNet-50 architecture was employed for model training. The trained model was then exported and used for real-time tracking via the DLC-Live method (18).

Tracking was performed by estimating the centroid of the mouse using the neck, back, and tail base key points, applied in both offline DeepLabCut and online DLC-Live tracking. For offline tracking, key points with a confidence score below 0.9 were interpolated to improve accuracy.

For offline freezing detection using DeepLabCut, the average movement of key points per frame was then calculated and analyzed alongside manually labeled freezing states. Using receiver operating characteristic (ROC) analysis, an optimal movement threshold was determined by identifying the point on the ROC curve with the minimum Euclidean distance *d* to the ideal point (where True Positive Rate = 1 and False Positive Rate = 0). This can be expressed mathematically as:

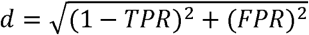

This threshold corresponding to the minimum *d* was selected as it maximizes classification accuracy for distinguishing freezing from non-freezing states. Frames with movement below this threshold for more than 0.5 seconds were classified as freezing states.

#### Accelerometry-based Offline Freezing Detection

Throughout the experiments, accelerometry data were continuously recorded at 30 kHz using an RHD2132 headstage equipped with a built-in 3-axis accelerometer (Intan Technologies LLC, Los Angeles). Data acquisition was performed using the Open Ephys system. The accelerometry data were calibrated, and a 1-Hz high-pass Butterworth filter (order 2) was applied to remove baseline drift. For each sample point, the acceleration vector magnitude *A_x_* was computed by combining the three directional components of the accelerometer data. The median absolute deviation (MAD) of *A_x_* was then calculated to establish a robust threshold for movement detection relative to baseline variability.

To detect freezing states, the absolute value of Ax was computed and smoothed to create a movement index. This index was segmented into continuous intervals by assessing threshold-crossing differences between adjacent movement indices. Specifically, segments were defined by determining if the difference between consecutive points exceeded a predefined threshold, allowing distance-based clustering of intervals with similar movement characteristics. Segments lasting longer than 0.5 seconds were retained for further analysis^21^.

To enhance the accuracy of freezing detection, threshold values were adjusted for freezing onset and offset. For freezing onset, a lower threshold (0.9 scaling) was applied, enabling sensitive detection of the initial drop in movement. The onset time of a potential freezing state was defined as the first time point at which the movement index fell below this adjusted threshold. For freezing offset, a higher threshold (1.1 scaling) was used, ensuring that freezing states were terminated only after a significant increase in movement. This bidirectional threshold adjustment allowed for sensitive initiation and robust termination of freezing detection.

### 2.5 Animals

The passive place avoidance task was conducted with 8 female mice. The social fear conditioning task involved 8 male experimental mice and 3 male stimulus mice, housed separately to prevent history of familiarization. All mice used in the experiments were adult C57BL/6J mice, aged 6–8 weeks old. They were housed in groups under standard conditions with a 12-hour reverse light/dark cycle (lights on from 19:30 to 7:30) and had free access to food and water. All experimental procedures complied with institutional animal care guidelines and received approval from the Animal Care and Use Committee at Fudan University Shanghai Medical College (Protocol number: DS-2021-165).

### 2.6 Behavioral task

#### Passive Place Avoidance Task

In the passive place avoidance task, mice were placed in a stationary, square-shaped shocker chamber, where they were expected to avoid entering a designated punishment area. This area, located in one corner of the chamber, covered one-fourth of the total surface and was associated with mild foot shocks.

The experiment began with a 20-minute free exploration (EXP) phase during which no shocks were delivered. RpiBeh was used to track the mice’s movement trajectories and record entries into the punishment area. For subsequent analysis, only the data from the first 10 minutes of this phase were used. After the exploration phase, the mice underwent two fear acquisition (ACQ) sessions, each lasting 10 minutes, with a 2-hour interval between sessions. During fear acquisition, mice received a foot shock whenever they entered the punishment area. The shock parameters were as follows: 0.7 mA intensity, 0.2-second delay after entering the area, 0.5-second shock duration, and a 2-second inter-shock interval (Coulbourn Instruments, ProBeCare, China).

On the second day, the mice were placed back in the shocker chamber for a 10-minute fear retrieval (RET) session. During this session, no electrical shocks were administered, allowing for the assessment of the mice’s fear memory for the punishment area and their avoidance behavior.

#### Social Fear Conditioning

This experiment lasted for two days. On the first day, the subject mice were placed in a three-chamber apparatus without any stimulus mice and allowed to explore freely for 10 minutes to familiarize themselves with the environment. After an interval, the mice underwent a three-chamber test.

On the second day, a conditioned stimulus procedure was conducted. The subject mouse was first placed in a shock chamber for 2 minutes of free exploration, during which it became briefly familiar with a stimulus mouse. The experiment then proceeded to a 15-minute conditioned stimulus phase under a closed-loop control system: whenever the subject mouse entered within a 5 cm radius of the stimulus mouse and stayed for more than 2 seconds, a 0.5-second foot shock of 0.7 mA was administered as a punitive stimulus. Three hours after the conditioned stimulus phase, the subject mice were tested again using the three-chamber apparatus.

Before the experiment began, the mice were housed in groups. Once the experiment started, however, each subject mouse was housed individually to prevent the extinction of fear memories through social interactions. Across the experiment, each subject mouse encountered three different stimulus mice, ensuring that all stimulus mice were novel. This setup aimed to eliminate the possibility of fear responses being directed at specific individuals. Additionally, the timing of both three-chamber tests was kept consistent for each subject to minimize the influence of circadian rhythms on social behavior, thereby reducing experimental variability.

To quantify the mouse’s social preference, a social preference index is calculated using the formula:

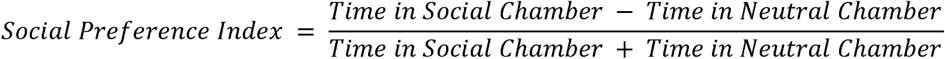

### 2.7 Statistics

Statistical analyses were performed using GraphPad Prism software. Behavioral data were analyzed through one-way ANOVA or nonparametric or mixed ANOVA, given the dataset’s characteristics. Post hoc multiple comparisons were performed using Dunnett’s or Tukey’s tests to assess differences between experimental groups. For comparisons between two conditions, paired t-tests were applied where relevant. Error bars in all graphs represent the standard error of the mean (SEM). Statistical significance thresholds were defined as *P < 0.05, **P < 0.01 and ****P < 0.0001.

## 3. Results

### 3.1 Position tracking and locomotion analysis

To assess the accuracy of position (center of mass, CoM) tracking, four different methods were compared: ANY-maze, DLC, DLC-Live, and Background Subtraction Method (BSM), with the workflow of the BSM algorithm shown in Fig.3A. Since CoM was defined differently in these methods (DLC/DLC-Live: centroid of neck, back, and tail base; BSM: CoM of largest component; ANY-maze: geometric center of the mouse’s region), minor discrepancies may naturally. Fig.3B. provides examples of 2D spatial coordinates, speed, and acceleration obtained from real-time detection by RpiBeh. Sample trajectory and heatmaps obtained with the BSM method are shown in Fig.3C. and Fig.3D.

**Fig. 3.**
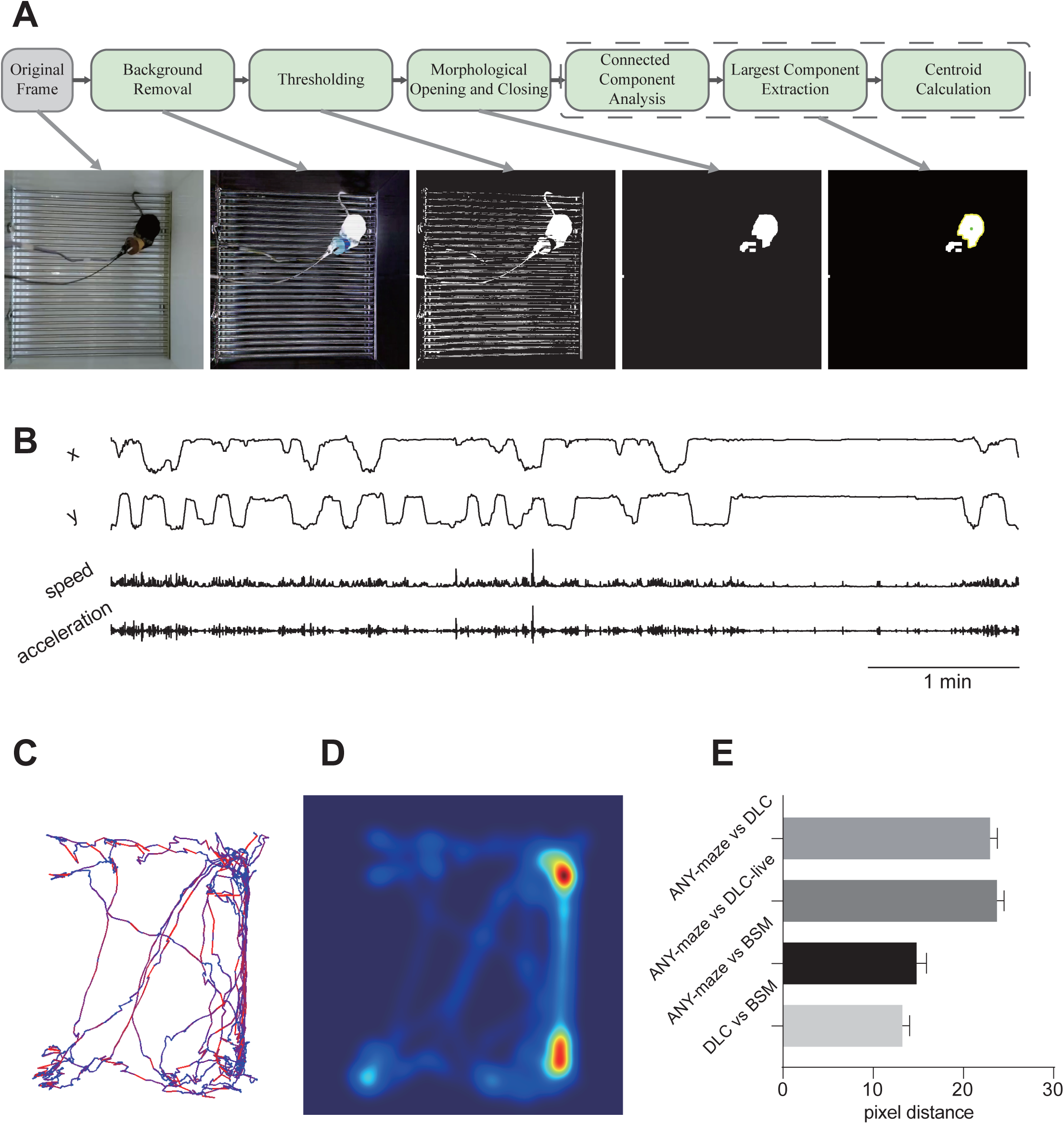
Performance of BSM based position tracking in Pavlovian fear conditioning. (A). Workflow of the background subtraction method. (B). Example of spatial coordinates (x: axis of abscissa; y: axis of ordinate; origin: top-left point), speed, and acceleration obtained from real-time detection by the RpiBeh. (C). Example of mouse trajectory (colors corresponds to speed change from slow (bluish) to fast (reddish)). (D). Example of a mouse position heatmap. (E). Distance between the center positions of the mouse detected using different algorithms (with 10 pixels approximately equivalent to 0.38 cm). Error bars represent the standard error of the mean (SEM).

The DLC method involved training a network to identify multiple body parts of the mouse.

The mouse’s center was calculated by taking the arithmetical mean of the coordinates of the neck, back, and tail base, with key points below 0.9 accuracy was linearly interpolated to ensure reliability. The DLC-Live method employed the same network model as DLC for real-time inference but did not apply interpolation, with the same selection of the neck, back, and tail base as the key points for center position calculation.

We computed the pixel distance between the center positions obtained by different methods for each frame and averaged these distances across each video. The comparison results are illustrated in Fig.3E. The results suggest that the BSM method yields center positions that are closely aligned with those from ANY-maze and DLC (10-15 pixels), with the distance between BSM and ANY-maze being smaller than that between ANY-maze and DLC. This result suggests that the BSM method effectively captures the mouse’s center position with high accuracy.

### 3.2 Online locomotion pattern detection

RpiBeh offers online detection and intervention capabilities for freezing, speed, and acceleration, enabling precise monitoring and control of locomotion patterns. Fig.4A. presents a sample frame non-matching ratio curve provided by RpiBeh, with the selected threshold set at 0.007. To validate the accuracy of fear detection, we analyzed 12 video recordings using multiple methods, including ANY-maze, Frame Difference, Offline Frame Difference (Offline FD), Accelerometer-Based Method (AUX), and the DeepLabCut (DLC) model. Manual annotations of fear states served as ground truth to benchmark the performance of these methods. Due to missing data in the AUX recordings, only 10 video sessions were analyzed. Fig.4B. shows intermediate processing steps of the Frame Difference (FD) algorithm. Fig.4C. illustrates the schematic of real-time speed pattern detection.

**Fig. 4.**
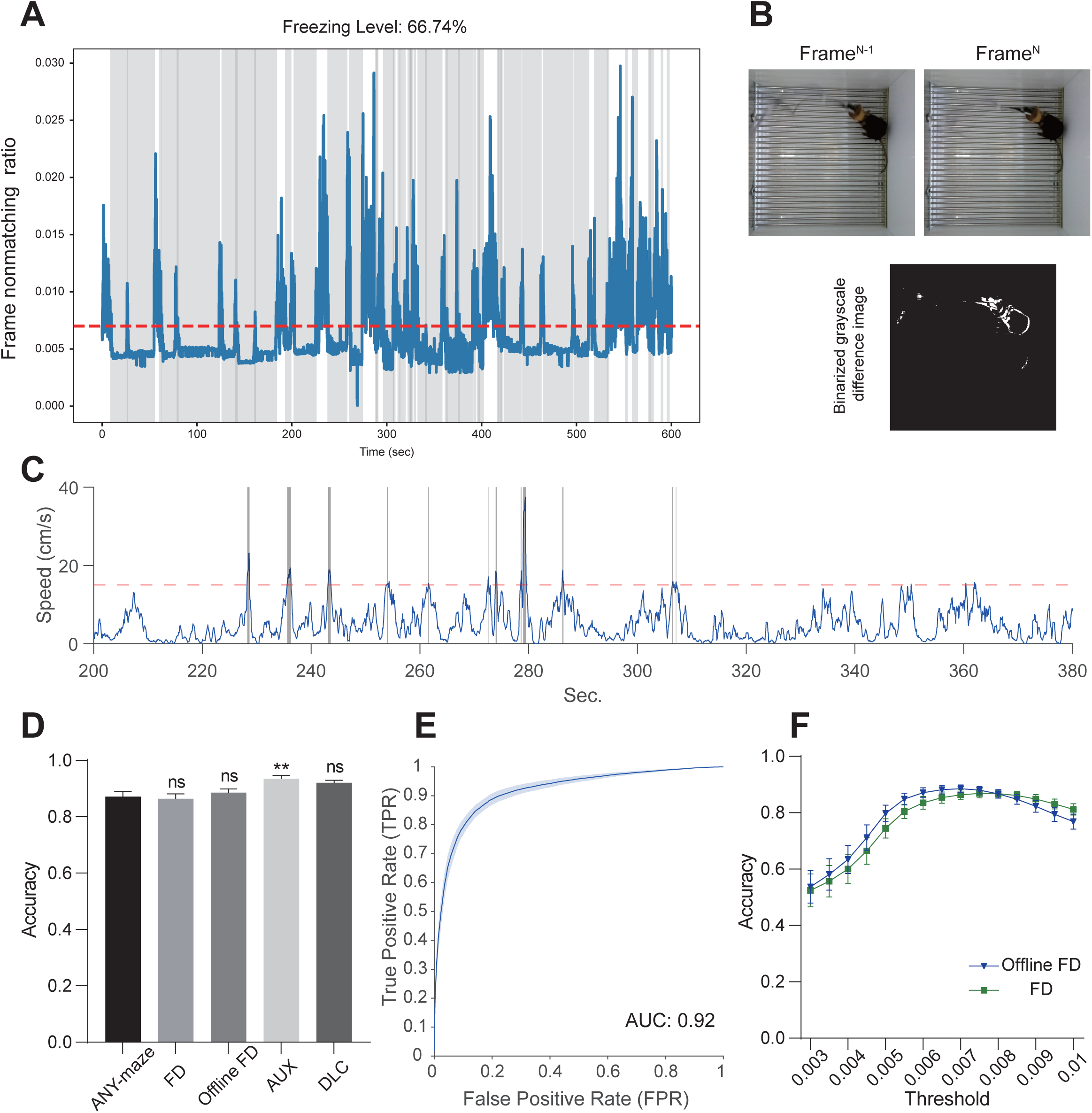
Performance of online locomotion pattern detection. (A). Schematic diagram of the online freezing analysis provided by RpiBeh (red dashed line indicates the selected threshold at 0.007). (B). Online speed detection schematic (gray area indicates speeds above the threshold, red line marks 15 cm/s). (C). Example images generated by the Frame Difference (FD) algorithm. (D). Comparison of freezing detection accuracy using different methods (FD, Offline FD with a threshold of 0.007). (E). ROC curve for detecting fear freezing status using frame nonmatching ratio. (F). Summary of FD algorithm accuracy in different threshold values. *P < 0.05; **P < 0.01; ****P < 0.0001; n = 12 (AUX: n = 10, due to missing data from two mice); One-way ANOVA (Nonparametric or Mixed ANOVA where applicable), followed by Dunnett’s multiple comparison post-hoc tests for C; Significancy indicators on the bars represent comparisons with the ANY-maze group for C. Error bars represent the standard error of the mean (SEM).

As shown in Fig.4D., the accuracy of FD, Offline FD, and DLC did not differ significantly from ANY-maze, indicating that FD offers reliable detection performance. Notably, the AUX method outperformed ANY-maze significantly, suggesting that when electrodes obstruct the animal’s visibility and electrophysiological recordings are involved, the AUX method provides more precise detection of mouse freezing states.

We further evaluated the classification performance of pixel difference ratio in distinguishing between fear freezing and non-freezing states. The ROC curve in Fig.4E. yielded an area under the curve (AUC) of 0.92, demonstrating the effectiveness of the pixel difference ratio in fear detection. Additionally, we investigated the impact of different threshold values on the accuracy of FD and Offline FD methods (Fig.4F.). The results show that accuracy first increases and then decreases with higher thresholds, indicating that the selection of the threshold is critical for optimal detection. As shown in Fig.4A., there is a distinguishable optimal range for threshold selection, enabling more accurate behavior classification.

### 3.3 Behavior-driven closed-loop intervention in PPA task

The experimental paradigm of the PPA Task is illustrated in Fig.5A. In this setup, a quarter of the arena is designated as a punishment zone, where mice receive mild electrical foot shocks upon entry. Initially, the mice are allowed to explore the arena freely without shocks (EXP phase). This is followed by two fear acquisition sessions (ACQ1 and ACQ2) and, on the second day, a fear memory retrieval session (RET).

**Fig. 5.**
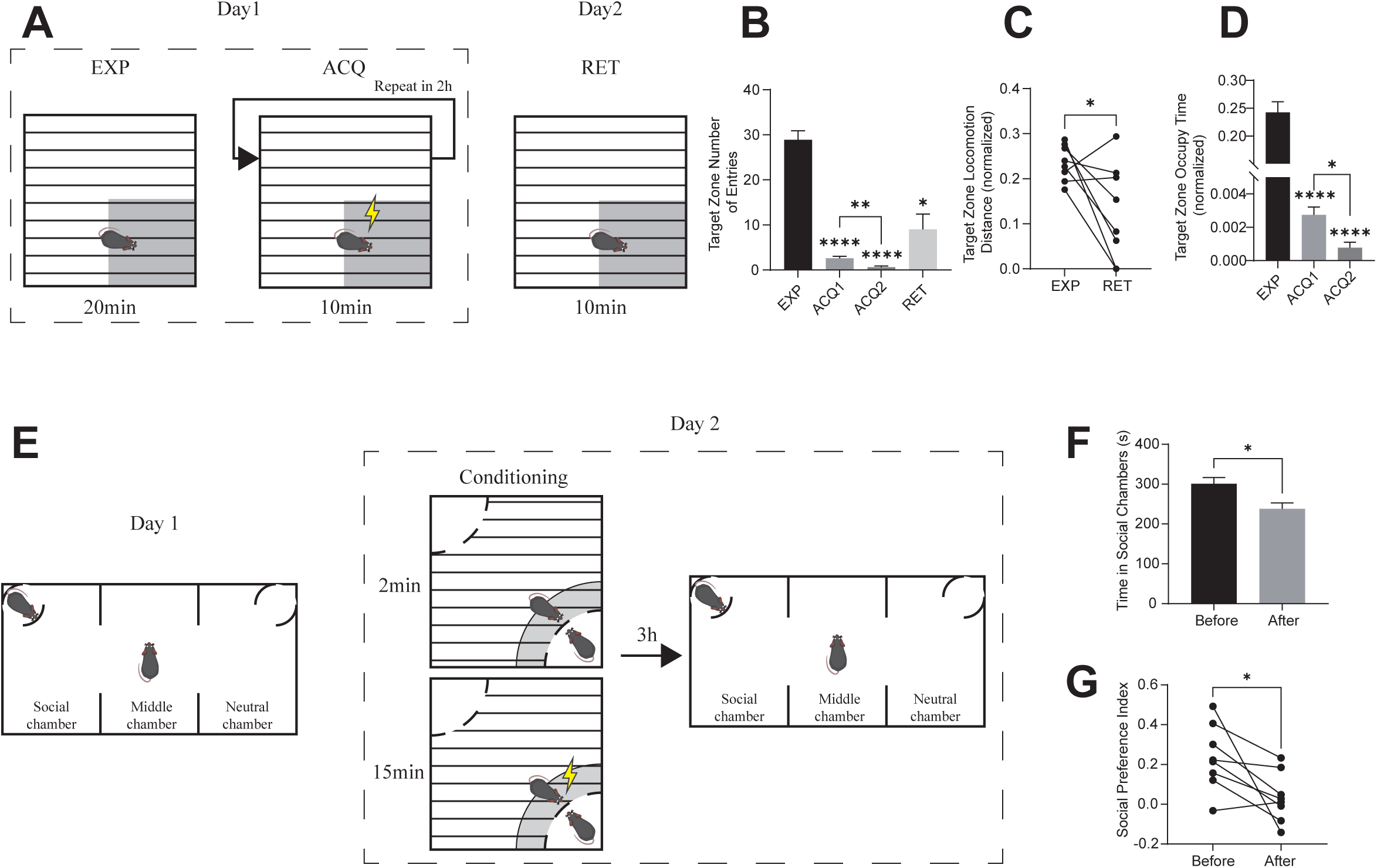
Behavioral-driven closed-loop experiments used RpiBeh. (A). Schematic diagram of the PPA Task. (B). Number of entries into the target (punishment) zone by the mice. (C). Ratio of distance traveled in the target (punishment) zone to the total distance, before and after stimulation (distance traveled within the target zone divided by the total distance). (D). Time spent in the target zone during fear acquisition (time spent in the target zone divided by the total session time). (E). Schematic diagram of the SFC Paradigm. (F). Time spent in the Social Chamber before and after conditioning. (G). Social preference index preceding and following conditioning, calculated as the difference in time spent between the Social Chamber and Neutral Chamber, normalized by the total time in both chambers, reflecting the preference for social interaction. *P < 0.05; **P < 0.01; ****P < 0.0001; n = 8; One-way ANOVA followed by Tukey’s multiple comparison post-hoc tests for B and D; Statistic significancy indicators on the bars represent comparisons with the EXP group; Paired t-tests for C, F, and G. Error bars represent the standard error of the mean (SEM).

As shown in Fig.5B., the number of entries into the punishment zone decreased significantly across the acquisition sessions, demonstrating successful learning of the avoidance behavior. On the second day, during the RET phase, the number of entries remained significantly lower than during the initial exploration (EXP), indicating memory retention of the learned avoidance behavior. Additionally, as depicted in Fig.5C. the ratio of distance traveled within the punishment zone to the total distance traveled during RET was significantly lower than that in the EXP phase, confirming that the mice actively avoided the punishment zone.

The analysis of stay duration further supports this observation (Fig.5D.) During the EXP phase, the stay time ratio (the proportion of time spent in the punishment zone relative to the total session time) was approximately 1/4, matching the proportional area of the punishment zone. This suggests that the mice initially exhibited no preference or avoidance for the punishment zone. However, as training progressed, the stay time ratio within the punishment zone decreased significantly, reflecting an acquired avoidance behavior. These results demonstrate that the software successfully enabled automatic closed-loop reinforcement of passive avoidance behavior in mice.

### 3.4 Behavior-driven closed-loop intervention in SFC

As shown in Fig.5E., mice SFC was established via ROI-based closed-loop intervention. In this paradigm, the experimental mouse first explores the shock box freely and briefly interacts with the social mate which was spatially restricted by a cylinder box. During the conditioning phase, RpiBeh performs real-time tracking of the experimental mouse’s position. When the experimental mouse approaches the stimulus mouse, it receives a mild foot shock during social interaction, which ultimately leading to the development of social avoidance behavior.

Before and after the conditioning session, the three-chamber test is conducted to evaluate the mouse’s social preference. As shown in Fig.5F., after the conditioning phase, the experimental mouse spent significantly less time in the Social Chamber. As depicted in Fig.4F., the social preference index also decreases significantly after conditioning, indicating reduced social motivation.

These findings demonstrate that the social fear conditioning paradigm was successfully established through ROI-triggered closed-loop intervention implemented by the RpiBeh, enabling precise manipulation of the mouse’s social behavior.

## 4. Discussion

In this study, we introduce RpiBeh, a Raspberry Pi and python language based multi-purpose open-source platform that integrates video recording, real-time position tracking, locomotion pattern analysis (speed, acceleration, immobility), and behavior-driven closed-loop intervention for rodent neuroethology. RpiBeh features a user-friendly graphic interface, good expandability for different behavior paradigms or closed-loop experiments, and currently supports the integration of DLC-Live, with potential for further expansion. In addition, RpiBeh equipped with frame-by-frame precision video synchronization. Those functionalities make RpiBeh an ideal choice for projects necessitates affordable and robust solutions for real-time processing and control of behavior data streaming in neuroscience.

One major advantage of RpiBeh established in its Raspberry Pi based video processing capacity, which has following merits: (1) frame-by-frame video synchronization was implemented by customizing the PiCamera library. Modern systems neuroscience experiments often demand parallel acquisition of multiple physiological and ethological signal streams. To achieve precise temporal synchronization over those data streams, many traditional solutions (e.g., Bonsai) depend on cost-ineffective industrial grade camaras that equipped with GIPO functionality. In contrary, RpiBeh adopted an software based strategy powered by Raspberry Pi ^5^, which sending out a train of brief TTL signals to allows post hoc alignment of behavioral and physiological events. (2) Compact and portable device., the Raspberry Pi camera used in the system is small (approximately 25 × 24 × 11.5 mm) and electrically isolated from AC power sources, minimizing potential electromagnetic interference. (3) Cost-effective and high accessibility. The hardware cost to implement all these functionalities is affordable for most research laboratories around the world.

RpiBeh had robust performance for application in various of different behavioral testing, such as position tracking in Pavlovian fear conditioning, spatial-ROI based automatic closed-loop reinforcement in PPA and SFC task, demonstrating the platform’s great potential for broad applications including learning and memory tasks, social behavior, locomotion and posture based behavioral analysis. Comparative test support that the our BSM algorithm and FD algorithm has accuracy as good as mainstream commercial software (e.g., Any-maze) or other popular deep learning empowered frameworks (e.g., DeepLabCut). In principle, the online locomotion pattern detection feature in RpiBeh allows for closed-loop applications based on custom defined behavioral features that beyond spatial ROI.

Importantly, we also demonstrated the utilization of DLC-Live for locomotion pattern detection within the RpiBeh framework, which could be expand into more sophisticated online posture estimation functionality ^8,22^. Last but least, RpiBeh also holds great potential for integrating with electrophysiological recording systems (e.g., Open Ephys) through enabling multi-modality (neurophysiological and behavioral) event driven closed-loop experiments and brain-computer interface applications, and promote which would be particularly valuable for applications like epilepsy detection ^23,24^, pain monitoring ^25–27^, and sleep intervention ^28,29^.

Despite its advantages, the software has certain limitations. Currently, it supports only single-animal tracking, although this limitation can be circumvented in specific paradigms. In the future, real-time multi-animal tracking could be achieved through the fusion of multiple video sources ^30^ or by leveraging deep neural networks ^11,12^. Similarly, while other third-party algorithms for behavioral analysis (e.g., DLC-Live) can be called in RpiBeh, many fine-grained built-in analysis features (e.g., posture detection) remain to be further developed.

## Acknowledgement

We thank all members of the Ni lab for discussion and support. This work is supported by National Key R&D Program of China (2024YFF1206500), National Science Foundation (T2394531), the Shanghai Municipal Science and Technology Major Project (2018SHZDZX01), ZJ Lab and Shanghai Center for Brain Science and Brain-Inspired Technology, China.

## Author Information

## Contributions

Project Design: Y.Sun, J.Zhang and J.Ni; Manuscript writing: Y.Sun, J.Zhang and J.Ni; Software implementation: Y.Sun, J.Zhang; Data analysis: Y.Sun, J.Zhang; Animal experiments: Y.Sun and Q.Wang; Project supervise and founding: J.Ni;

## Data Availability

The datasets used and/or analyzed during the current study are available from the corresponding author upon request. At the time of publication, RpiBeh is available at: https://github.com/NiLab-FDU/RpiBeh

